# Concerted interactions between multiple gp41 trimers and the target cell lipidome may be required for HIV-1 entry

**DOI:** 10.1101/2020.04.18.048173

**Authors:** Biswajit Gorai, Anil Kumar Sahoo, Anand Srivastava, Narendra M. Dixit, Prabal K. Maiti

## Abstract

The HIV-1 envelope glycoprotein gp41 mediates the fusion between viral and host cell membranes leading to virus entry and target cell infection. Despite years of research, important aspects of this process such as the number of gp41 trimers involved and how they orchestrate the rearrangement of the lipids in the apposed membranes along the fusion pathway remain obscure. To elucidate these molecular underpinnings, we performed coarse-grained molecular dynamics simulations of HIV-1 virions pinned to the CD4 T cell membrane by different numbers of gp41 trimers. We built realistic cell and viral membranes by mimicking their respective lipid compositions. We found that a single gp41 was inadequate for mediating fusion. Lipid mixing between membranes, indicating the onset of fusion, was efficient when 3 or more gp41 trimers pinned the membranes. The gp41 trimers interacted strongly with many different lipids in the host cell membrane, triggering lipid configurational rearrangements, exchange, and mixing. Simpler membranes, comprising fewer lipid types, displayed strong resistance to fusion, revealing the crucial role of the lipidomes in HIV-1 entry. Performing simulations at different temperatures, we estimated the free energy barrier to lipid mixing, and hence membrane stalk formation, with 4 tethering gp41 trimers to be ~6.2 kcal/mol, a >4-fold reduction over estimates without gp41. Together, these findings present molecular-level, quantitative insights into the early stages of gp41-mediated HIV-1 entry. Preventing the requisite gp41 molecules from tethering the membranes or altering membrane lipid compositions may be potential intervention strategies.

**SIGNIFICANCE:** Interactions between viral envelope proteins and host cell surface receptors leading to HIV-1 entry are well studied, however the role of membrane lipids remains obscure, although entry hinges on lipid mixing and the fusion of viral and cell membranes. We performed detailed simulations of HIV-1 and target cell membranes tethered by viral gp41 trimeric proteins to elucidate the proteo-lipidic contributions to viral entry. We found that the cooperative effects of multiple gp41 trimers and natural lipidomes of the membranes facilitate membrane fusion. The functional domains of gp41 altered local lipid concentrations, reduced membrane repulsions, and facilitated inter-membrane lipid mixing. These molecular-level insights offer a glimpse of the cryptic mechanisms underlying HIV-1 entry and suggest new interventions to combat HIV-1 infection.

## 1 INTRODUCTION

The entry of HIV-1 into its target CD4 T cells is mediated by its surface envelope protein, Env, a homotrimer of noncovalently linked heterodimers of glycoproteins gp120 and gp41 (1–4). Entry begins with gp120 binding to the cell surface receptor CD4, following which conformational changes in gp120 enable its binding to one of the co-receptors CCR5 or CXCR4. Further conformational changes ensue, resulting in the exposure of the N-terminal fusogenic domain of gp41, also called the fusion peptide (FP; see Fig. S1), its insertion into the target cell membrane, and membrane fusion leading to virus entry (2). Blocking virus entry is an important intervention strategy. Drugs that block CCR5 binding to gp120 (5) or conformational changes in gp41 facilitating entry (6) are approved for clinical use. Vaccination strategies also aim to administer or elicit antibodies that target Env and prevent entry (7). The success of these strategies relies on a detailed, quantitative understanding of the molecular requirements of the entry process, which despite years of research, is still lacking. For instance, how many Env trimers are required for entry is yet to be established (8).

While attention has been focused in previous studies on the proteins involved in the entry process (4, 8-10), much less is known of the roles played by the lipids involved despite inter-membrane lipid mixing being central to membrane fusion. Lipid mixing may be an important barrier to fusion given that the lipidomes of the HIV-1 and target cell membranes are vastly different (11). The proteins and lipids must thus work in tandem to orchestrate virus entry.

Here, to elucidate the roles of both the lipids and the associated proteins in HIV-1 entry, we performed detailed, long-timescale (~10 μs), coarse-grained molecular dynamics (CGMD) simulations of the viral and host cell membranes held in close apposition by gp41 trimeric tethers. We faithfully mimicked the known compositions of the viral and host lipidomes, leading to the creation of complex, multi-lipid-type membranes. By varying the number of gp41 tethers, we estimated the protein requirements for lipid mixing and their energetic contributions. At the same time, we unravelled the detailed configurational changes of the lipids culminating in mixing and eventually membrane fusion. Specifically, we answered the following questions: (i) How many gp41 trimers are required for membrane fusion? (ii) How do they orchestrate lipid mixing? (iii) What is the associated free energy barrier? (iv) How important are the lipid compositions to membrane fusion?

## 2 RESULTS

### More than 2 gp41 trimeric units are necessary for fusion

Previous studies, using experiments and simulations, have demonstrated the possibility of membrane fusion without the involvement of any fusogenic peptides or proteins (12–14). We therefore first performed simulations containing the HIV-1 and human T cell membrane models in the absence of gp41. The membranes differed in their overall lipid compositions as well as in the compositions of their inner and outer leaflets (Methods). We built HIV-1 as a vesicle of 20 nm diameter and held it initially in close proximity to a flat 40 × 40 nm^2^ cell membrane (Fig. 1A), representative of the virus-host cell membrane interactions considering the cell diameter is nearly 100-fold the virion diameter. Rendering the cell membrane as a vesicle of 20 nm diameter did not alter our findings (Fig. S2). We found that the membranes drifted away from each other in our 5 *μ*s simulations (Figs. 1A and S2). A previous study with vesicles composed of 1-2 lipid types found them repelling during CGMD simulation (15). The electrostatic repulsion due to the polar lipid head groups and the hydration force seems to drive the membranes apart (16). Within the duration of our simulations, thus, the membranes were resistant to spontaneous fusion. We, therefore, examined next whether gp41 trimers could overcome this resistance.

**Fig. 1.**
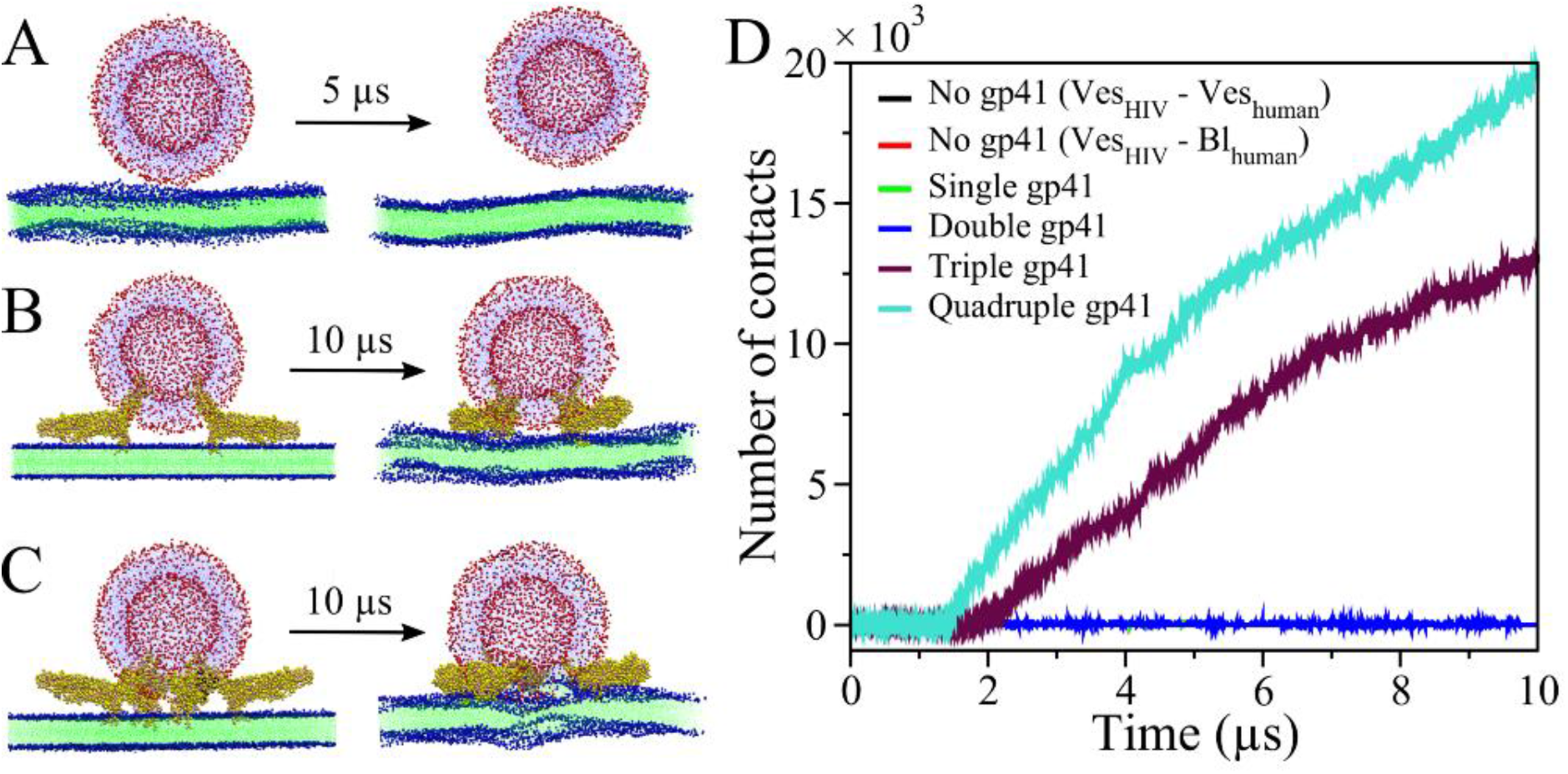
Stoichiometry of gp41 for membrane fusion. The instantaneous snapshots of the initial (left) and final (right) configurations of the system for fusion of the human bilayer–HIV-1 vesicle in the presence of (*A*) no, (*B*) double, and (*C*) quadruple gp41 trimeric units are depicted. (*D*) The number of contacts among inter-membrane lipid beads are shown for the simulated systems at 300 K with different gp41 trimeric units (0–4), which form bridges between the human bilayer and the HIV-1 vesicle membrane models. Note that black, red, and green color lines are always zero, hence not visible in the plot. The exchange of lipids between the human bilayer and the HIV-1 vesicle is observed only in the presence of three and four gp41 trimers. The lipid head groups and tails of the human bilayer are shown in vdW (blue) and line (green) representations, respectively. The lipid head groups and tails of the HIV-1 vesicle are shown in vdW (red) and line (light blue) representations, respectively. The gp41 trimeric unit is shown in vdW representation with a yellow color. Water and ions are not shown here for clarity.

We now initialized our simulations with the membranes apposed as above but tethered to each other by different numbers of gp41 trimers (Figs. 1B, C, and S3). The gp41 trimers were placed symmetrically around a focal region. We found that with 1 and 2 gp41 trimers, the membranes remained apposed but formed no inter-membrane lipid contacts in the 10 *μ*s simulations (Fig. 1D). Extending the simulation with 2 gp41 trimers from 10 *μ*s to 20 *μ*s did not change the scenario (Fig. S4). With 3 gp41 trimers, the number of contacts increased rapidly starting at 1.8 *μ*s, indicating the onset of lipid mixing. With 4 trimers, mixing began earlier, at 1.4 *μ*s, and proceeded faster. These simulations demonstrate that, for the initial gp41 configurations chosen, at least 3 trimers were necessary to initiate fusion. This finding is consistent with recent experiments showing that most HIV-1 strains require 2-3 trimers for entry (8). Further, fusion was expedited with more trimers involved.

### gp41 trimers act cooperatively to initiate membrane fusion

The multiple gp41 trimers involved in fusion initiation could act independently or cooperatively. To ascertain this, we tracked the movements of the trimers throughout the simulations. Specifically, we estimated the distances between the center-of-mass of every pair of gp41 trimers, considering their fusion peptides and transmembrane domains. We numbered the trimeric units, U1, U2, etc. We found that the gp41 units were mobile throughout the 10 *μ*s long simulation and had a tendency to cluster transiently (Fig. 2). For the system with 3 gp41 trimers, U1 and U3, which were initially separated by ~9 nm, approached each other, reached a minimum separation of ~3.5 nm at ~2 *μ*s, and then moved away (Fig. 2A, C). Interestingly, the time at minimum separation corresponded to the time when inter-membrane contacts began to increase rapidly (Fig. 1D). With the 4 gp41 trimer system too, U1 and U2 achieved a minimum separation similar to that above and at a time, ~1.5 *μ*s, that corresponded well with the rise of inter-membrane contacts (Figs. 2D and 1D). The other pairs did not come this close. It appears thus that when two gp41 trimers approach each other within a separation of ~3.5 nm, they can trigger the formation of lipid contacts and initiate membrane fusion. The additional trimers perhaps serve to stabilize the membranes when lipid mixing is ongoing. It is possible that in our simulations with 2 gp41 trimers, because the trimers were placed diametrically opposite each other, they never approached such close separations within our simulation time. Env spikes of a mature HIV-1 virion have been observed to form small clusters (17), possibly to drive fusion. To the best of our knowledge, we report for the first time the minimum distance of approach between two gp41 trimeric units that is essential to initiate inter-membrane lipid contact and hence fusion. This observation may have implications for vaccines aimed at neutralizing HIV-1 virions (7). We turn next to the role of lipids.

**Fig. 2.**
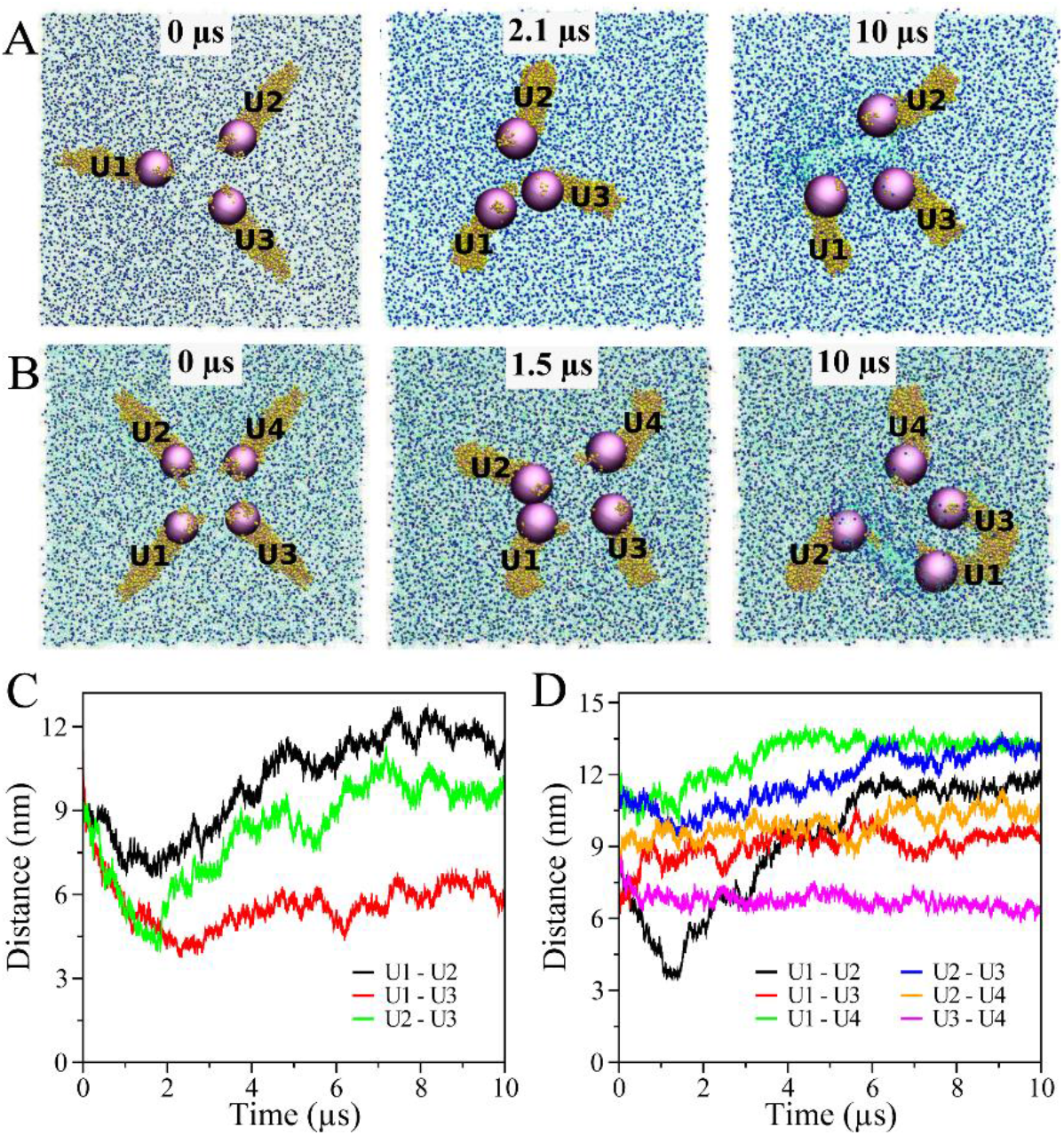
Cooperative action of gp41 trimer units (U) to initiate lipid mixing and membrane fusion. The snapshots at the initial, onset of stalk formation, and final simulation time instances for systems with (*A*) triple (U1-U3) and (*B*) quadruple (U1–U4) gp41 units are depicted. The gp41 trimeric units and the center-of-mass (CoM) of the FP+TMD domains of each gp41 trimer are represented by yellow beads and violet sphere, respectively. The hydrophobic tails and the polar heads of the human bilayer lipids are represented by green lines and blue vdW spheres, respectively. The simulation time is inscribed on the top-middle of each figure. The HIV-1 vesicle, water beads, and ions are not shown in the snapshots for clarity. For each pair of gp41 trimeric units, the distance (in the *X–Y* plane) between the CoMs of the FP+TMD domains as a function of the simulation time is shown for systems with (*C*) triple and (*D*) quadruple gp41 units. The gp41 trimeric units seems mobile in nature and approach each other (with the minimum possible separation of ~3.5 nm) to initiate the stalk formation, which facilitates inter-membrane lipid mixing. After the stalk formation process completes, the separations between gp41 trimer units escalate to further proliferate the stalk elongation.

### Viral and cell lipidomes promote membrane fusion

Cell membranes are composed of several lipid types. Further, the lipid compositions of the inner and outer leaflets of membranes can be quite different. Most computational and experimental studies consider simple membrane models composed of only one or at best a few lipid types (18). To examine whether such simple models capture the HIV-1 fusion process, we performed simulations with the HIV-1 vesicle and the apposed cell membrane both composed of POPC (80%) and POPS (20%) CG lipids, tethered by 4 gp41 trimers (Fig. S5). We found no inter-membrane contact formation within the 10 *μ*s long simulations performed at the regular temperature (300 K) or an elevated temperature (400 K). As we showed above, however, membrane models composed of near biological lipid compositions formed lipid contacts and mixed (Fig. 1D). The lipidomes of the virus and cell membranes are thus important to membrane fusion; they appear to accelerate fusion. We examined next how the lipidomes participated in the fusion process.

### Differential splaying of lipid types may promote fusion

Membrane fusion involves changes in membrane lipid configurations as well as mixing and homogenization of lipid compositions. At the local level, configurational changes may manifest as tilt and splay. We examined both and compared them between lipids near the focal, contact region, within a 20 Å distance, and those far away. Initially, the lipids exhibited no tilting and were aligned along the bilayer normal (Fig. S6). At the onset of lipid mixing, substantial tilt away from the normal was seen, in keeping with the bending membrane (Fig. S6). Further, different lipid types showed different extents of tail splay in the process, quantified as the probability distribution of the splay angle between the aliphatic tails (Fig. S7). In the absence of fusion proteins, splaying of lipid tails facilitates membrane fusion (19). Away from the contact region, the lipids were unstressed and exhibited splay within a narrow range, 35°–42°. In the contact region, the splay varied far more, 20°–60°. DOPS, with one double bond in both the aliphatic tails, showed the highest splay. The splay angle of a few DOPS and POPS molecules, which are found mostly in the inner leaflets, exceeded 60°, consistent with the greater splay of inner leaflet lipids observed with the DMPC/DOPE membrane system (20). These observations suggest that some lipids splay more than others and absorb more of the stress associated with membrane bending. Other lipids that are less flexible can get away with much less splay, and perhaps even contribute to membrane integrity. In a model membrane lacking this lipidome diversity, the lipids may either all splay, leading to poorer integrity, or not splay at all, preventing fusion. The lipidome diversity may thus serve as an optimum in this integrity-flexibility trade-off.

### Lipid transfer between membranes is asymmetric and asynchronous

To quantify lipid mixing between the membranes, we tracked the number of lipids transferred from the cell membrane to the HIV-1 vesicle (C→H) and the same from the HIV-1 vesicle to the cell membrane (H→C) for each lipid type. For each lipid type, C→H transfer was observed immediately after the lipid contact initiation time (~1.4 *μ*s), whereas H→C transfer occurred much later, ~4 *μ*s (Fig. 3). Also, the C→H transfers (in 10 *μ*s) were fewer than H→C transfers. The cell membrane underwent significant morphological changes during the process, whereas the HIV-1 membrane maintained its spherical geometry. The HIV-1 membrane contains more cholesterol (~45%) than the cell membrane (~37%), which provides it more order and rigidity, possibly explaining the fewer successful lipid transfers to the vesicle than from it. In descending order of numbers, C→H transfers followed: Cholesterol > POPC > POSM ≈ DPPC > POPE ≈ DPPE; whereas H→C transfers followed: Cholesterol > POPE ≈ POSM > POPC ≈ POPS. As cholesterol has the ability to flip-flop between the inner-outer leaflets (Fig. S8) and induce negative curvature to the membrane (21), it shows the highest tendency to translocate between the membranes in the focal region. In our simulations, other lipid types did not flip even once within 10 *μ*s (Fig. S8); transfers were thus predominantly of the lipids in the outer leaflets. For all lipid types except cholesterol, the transfers saturated after 8 *μ*s, indicative of the achievement of new stable compositions. (Fig. 3).

**Fig. 3.**
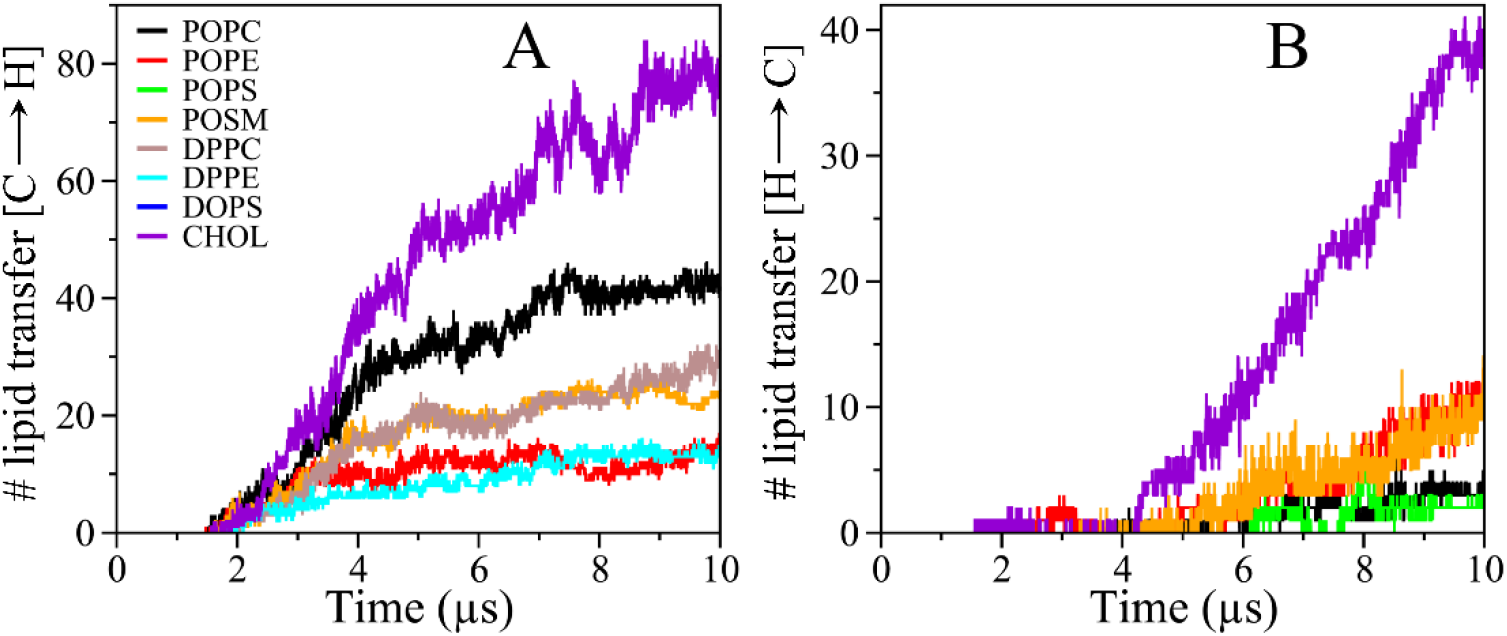
Kinetics of inter-membrane lipid transfer. For each lipid type, the number of lipids transferred from the human membrane bilayer to the HIV-1 vesicle (*A*) and from the HIV-1 vesicle to the human membrane bilayer (*B*) is plotted as a function of the simulation time. The human membrane contains every lipid types except POPS, while the HIV-1 vesicle contains only POPC, POPE, POPS, POSM, and cholesterol.

### gp41 domains trigger local alterations in lipid compositions

Given that no lipid mixing occurred in the absence of gp41 tethers, we examined the interactions between gp41 and the cell membrane lipids, which could potentially have facilitated the mixing. We noticed a marginal elevation in the concentrations of POPC, POPE, POSM, and cholesterol in the focal region (Fig. S9). The interactions of functional domains of gp41 with lipid membranes have been studied using different biochemical and biophysical techniques (22). However, their preferences for lipid types and their roles in membrane fusion remain unclear. We calculated the 2-dimensional radial distribution functions (RDF) of the various membrane components around each gp41 functional domain and used the heights of the first peaks in the RDFs to infer the lipid association propensities. The resulting preferences are summarized in Fig. 4.

**Fig. 4.**
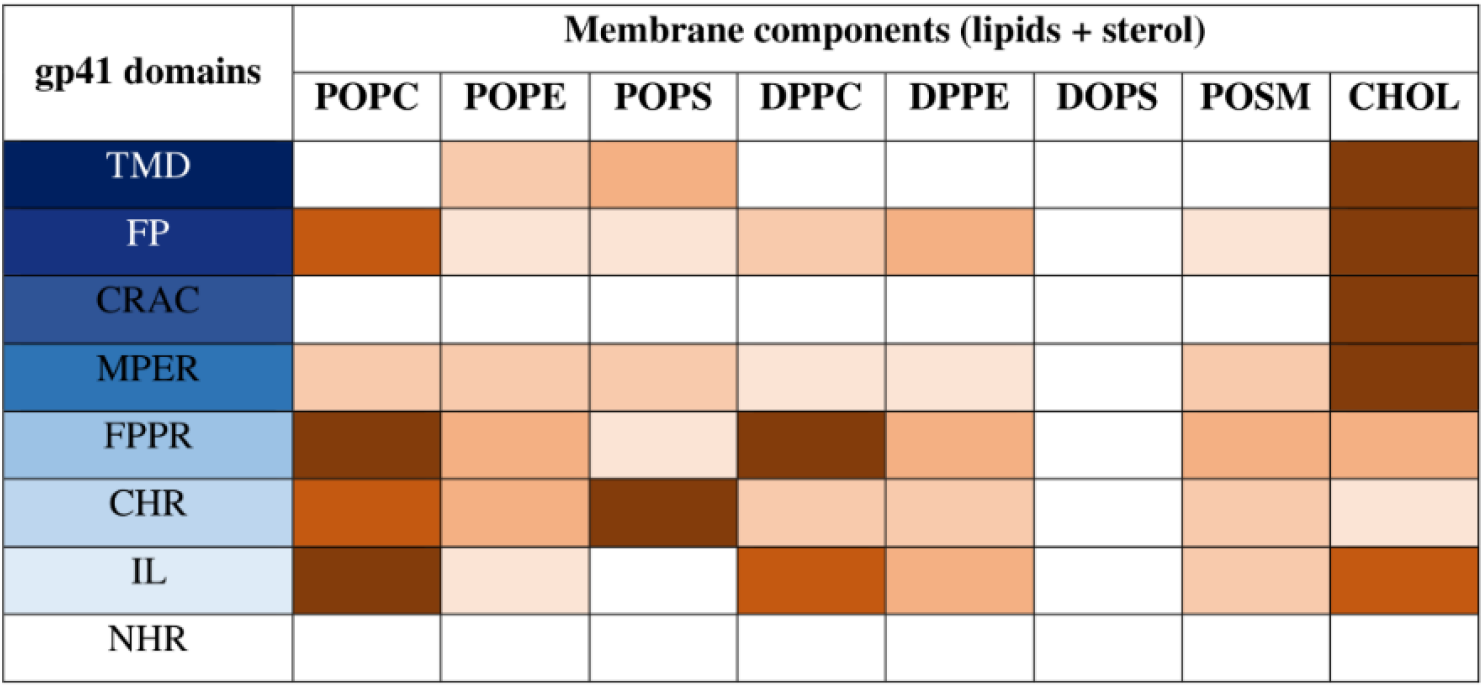
Summary of the interactions of various gp41 functional domains with the different lipid types. The occupancy of each lipid type around the functional domains of gp41 are depicted in the tabular form. The likeliness of each gp41 functional domain (when present in the six-helix bundled conformer) for the lipids varies from strong to weak, which is represented by the dark blue-to-white color gradient. TMD and FP domains of gp41, initially placed in the HIV-1 and human membranes, respectively, show the maximal fondness for the lipids, which is followed by CRAC, MPER, FPPR, CHR, and IL. Note that there is no interaction between NHR and the lipids, as NHR is enclosed by the CHR helices in the 6HB post-fusion conformer of the gp41 trimer (Fig. S1). The relative occupancy of each lipid type around the functional gp41 domains varies from strong to weak, which are represented in the dark brown-to-white color gradient. As the anionic DOPS lipid is only present in the inner leaflet of the human bilayer, it does not interact with any functional domain of gp41. All other lipid types are found to interact with at least one of the functional domains of gp41.

The transmembrane domain (TMD) showed the highest preference for cholesterol, followed by the negatively charged POPS and the zwitterionic POPE lipids. The fusion peptide (FP) too showed the highest preference for cholesterol followed by zwitterionic lipids. It, however, showed minimal affinity for the negatively charged lipids. The membrane-proximal external region (MPER) too showed the highest preference for cholesterol, followed by zwitterionic and anionic lipids. The later is argued to be due to MPER’s cholesterol recognition amino acid consensus (CRAC) motif found at the C-terminal end (23). RDFs for the CRAC motif also showed that only cholesterol was present near it. Note that MPER is considered as the functional counterpart of FP and plays crucial role in bridging HIV-1 and host membrane (22). The fusion peptide proximal region (FPPR) domain was surrounded by zwitterionic lipids more than anionic lipids or cholesterol. The C-terminal heptad repeat (CHR) preferred the negatively charged POPS the most, followed by the unsaturated zwitterionic lipids. On the other hand, the immunodominant loop (IL) showed higher affinities for zwitterionic lipids followed by cholesterol.

A recent experimental study has argued that cholesterol-rich domains facilitate gp41-mediated membrane fusion (24). Our simulations showed the preferred localization of cholesterol around FP, CRAC, and MPER domains of gp41. This localization, given the hydrophobic nature of cholesterol, would reduce the unfavorable interactions between the apposed membranes near the gp41 tethers. When a sufficient number of gp41 trimers accumulated, the opposing interactions were sufficiently reduced to allow lipid transfers and mixing. Our study thus reveal the crucial role of FP, CRAC, and MPER domains in membrane stalk formation. They cause local redistributions of the lipid types, enriching cholesterol, thereby mitigating membrane repulsions and facilitating fusion.

The significance of the IL domain during fusion is not well understood. We find from the simulations that IL anchors into the cell membrane and induces membrane bending. Other domains seem to play an insignificant role in gp41-mediated membrane fusion.

### gp41 reduces the activation barrier for membrane stalk formation

The free-energy landscape for HIV-1–cell membrane fusion is complex (Fig. 5A), as it encompasses several intermediate processes including the formation of the membrane stalk, the hemifusion diaphragm, and the final fusion pore (25). It is known from various studies that the free-energy barrier for the initial stalk formation, in the absence of proteins, is quite high (~40–220 *kT*) (26), mainly due to the electrostatic repulsion between the polar head groups and the hydration repulsion. Our simulations showed that gp41 trimers facilitate lipid mixing, leading to stalk formation. To quantify the extent to which gp41 trimers reduce the free-energy barrier (ΔG_*S*_), we performed simulations with 4 gp41 trimers tethering the membranes (Fig. 1C) at 5 different temperatures in the range 300–400 K. We found that the time of onset of lipid contact formation, or the stalk initiation time, *τ*, decreased with the temperature, *T.* This implied that stalk formation is an activated process, as depicted in Fig. 5A. We used the temperature dependence of *τ* to estimate ΔG_*S*_ using the Arrhenius equation, *τ* = *τ*_0_ *exp*(ΔG_*S*_/*kBT*), where *τ*_0_ is the intrinsic time constant and *kB* is Boltzmann’s constant. We found that ΔG_*S*_ was 6.2 ± 1.0 kcal/mol, or ~ 10.5 ± 1.7 *kT.* This is a reduction of >4-fold compared to the barrier without fusion proteins (26), indicating the critical role played by gp41 in facilitating HIV-1 entry. Because lipid mixing is expedited with increasing gp41 trimers (Fig. 1D), it follows that ΔG_*S*_ may be reduced further with more gp41 trimers.

**Fig. 5.**
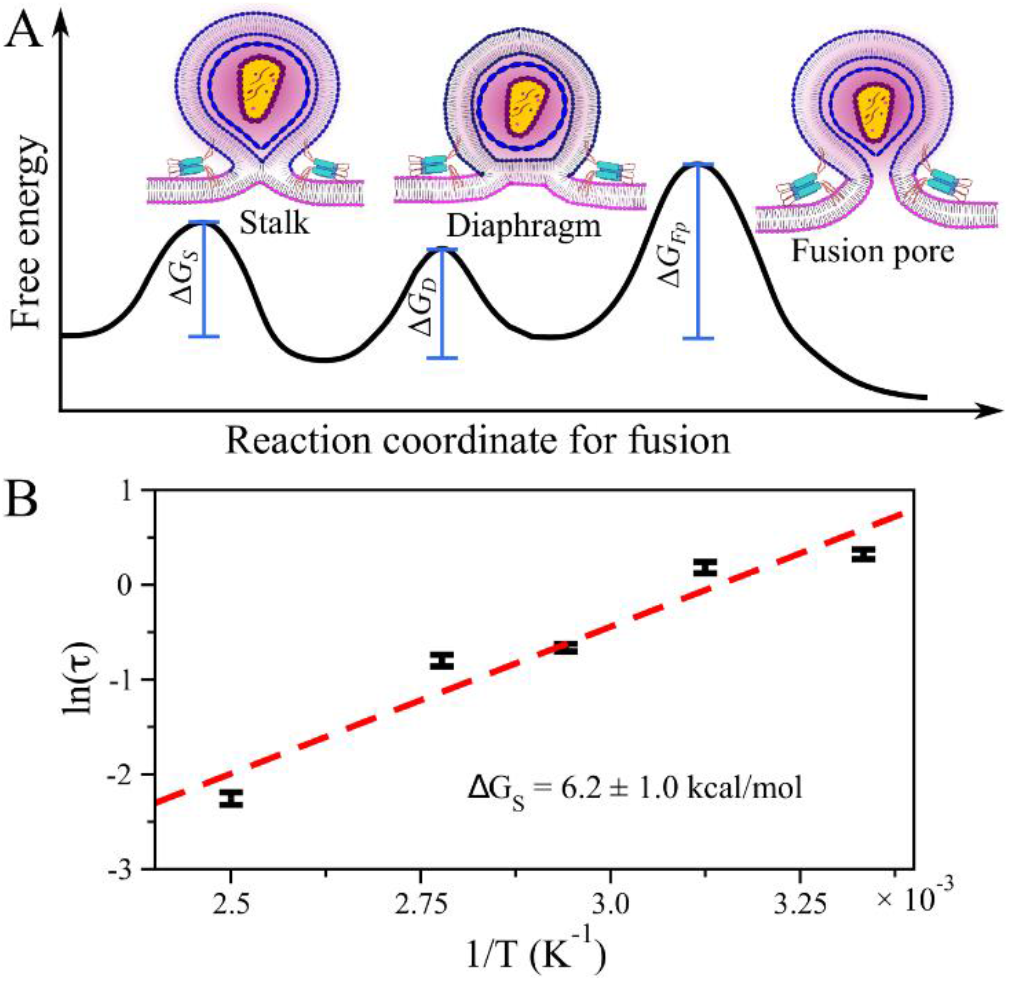
The activation barrier for the gp41-mediated stalk formation. (*A*) Schematic of the free-energy landscape for the human-HIV-1 cell membranes fusion, comprising of the activation barriers for the initial stalk formation (ΔG_*S*_), the intermediate stalk elongation or hemifusion diaphragm (ΔG_*D*_), and final fusion pore formation (ΔG_*f_p_*_), is shown. Each of the three energy-driven processes along the fusion pathway is inscribed in the schematic. The human cell is shown as a membrane bilayer with magenta color lipid heads, whereas the HIV-1 virion, enclosing the capsid, is depicted as a vesicle with blue color lipid heads. The gp41 trimeric unit is shown in cyan color. (*B*) The natural logarithm of the stalk initiation time (*τ*) is plotted as a function of the inverse temperature. The five data points (from right to left) correspond to temperatures 300 K, 320 K, 340 K, 360 K, and 400 K, respectively. The error bar is the standard deviation of the mean *τ* estimated from two independent simulations at each temperature. ΔG_*S*_ (= 6.2 ± 1.0 kcal/mol) is obtained by linear fit (dashed line) of the data points to Arrhenius formula, ln *τ* = ln *τ*_0_ + Δ*G_S_*/*kBT*.

## 3 DISCUSSION

Targeting HIV-1 entry into cells is an attractive intervention strategy and is central to ongoing efforts to design potent HIV-1 vaccines (7). In this study, we elucidate important molecular-level details governing the initiation of membrane fusion following the anchoring of gp41 trimers into the target cell membrane. Using unbiased CGMD simulations, we show that the concerted action of multiple gp41 trimers and membrane lipidomes are required for membrane fusion. When 3 or more gp41 trimers tethered the membranes, two gp41 trimers approached each other to within a certain distance, induced local changes in the lipid composition, which reduced the repulsive barriers between the apposed membranes and facilitated lipid transfer across the membranes leading to membrane fusion (Fig. S10). It follows that interventions that prevent the requisite number of gp41 trimers from tethering the membranes, prevent the approach of trimer pairs to within a certain distance, or alter the lipidomes may all present new avenues to block HIV-1 entry.

Previous studies have inferred that the number of fusogenic proteins required for HIV-1 entry varies from 1–8, with most HIV-1 strains requiring 2-3 (8, 27). In accordance, our study shows that at least 3 gp41 trimers were necessary for membrane fusion initiation. Further, the fusion protein-induced reduction of the free energy barrier to membrane fusion we estimated is in keeping with recent experimental and theoretical studies (8, 28, 29). We find that the barrier reduces with the number of gp41 trimers involved and that with 4 gp41 trimers, the barrier is reduced >4-fold over that in the absence of the trimers. Antibodies that target gp41, especially the MPER region, have been identified, can neutralize a wide variety of HIV-1 strains (30).

Our study, like the previous ones (8), provides a quantitative guideline for the use of such antibodies. The clustering of the Env proteins observed near the virus-cell contact region (4) implies that nearly all of the Env trimers may be involved in tethering the membranes. Cryo-electron microscopy has revealed that HIV-1 virions carry a mean of 14 Env trimers (31). Thus, antibodies must block 12 or more Env trimers on average to prevent HIV-1 entry. These considerations may help define targets for antibody elicitation using vaccination or the administration of such antibodies using passive immunization strategies. The observation that gp41 trimers move to within ~3.5 nm at the point of lipid contact formation across membranes suggests that preventing such apposition may prevent HIV-1 entry. One can imagine antibodies or other agents that can bind gp41 and occlude a region of ~3.5 nm around the gp41 on the cell membrane. Such agents may act as novel HIV-1 entry inhibitors. Our study, for the first time, elucidates the important role of the membrane lipidomes in HIV-1 entry. The lipidomes of HIV-1 and host cell membranes comprise many different lipid types (11). Simpler membranes that use fewer lipid types did not lead to successful membrane fusion initiation in our simulations. Two major effects allowed biologically realistic membrane lipidomes to facilitate fusion. First, membrane bending saw some lipids splay more while others less, the former absorbing much of the bending stress and conferring flexibility to the membrane while the later potentially maintaining membrane integrity. Membranes with fewer lipid types may not achieve this apportioning required to strike a balance between flexibility and integrity, rendering fusion difficult. Second, the lipidomes interacted with gp41 functional domains, which altered local lipid compositions in a way that reduced the repulsive forces between the membranes. Specifically, we found prominent relocalization of cholesterol near gp41, which because of its hydrophobic nature reduced hydration repulsions. Importantly, when multiple gp41 trimers were closely positioned, the effect reached a threshold that allowed lipid contact formation and mixing. Thus, the concerted activity of multiple gp41 trimers and the lipidomes were involved. Interventions could consider altering the lipidomes, which may prevent such lipid reorganization and preclude fusion. Indeed, altering membrane cholesterol has already been shown to compromise HIV-1 fusion (32).

Our simulations have focussed on the narrow window in the HIV-1 entry process between gp41 anchoring and lipid mixing. This window, however, is a crucial one and requires difficult simulations involving both membrane structures, viral structures, lipids, and proteins. The events preceding gp41 anchoring are extracellular and can be studied without explicit membrane constructions. The events post lipid mixing relies less on the role of proteins and lipidome diversity and thus can be studied using simpler model membranes without proteins. Our study, thus, addresses the most challenging part of the entry process. Indeed, we are thus able to unravel the crucial roles of the lipidomes and the cooperative functions of multiple gp41 trimers. Future studies may integrate our findings into larger-scale simulations starting with gp120-CD4 binding and culminating in fusion pore formation, which may soon become tractable as quantum computing technologies mature.

## ACKNOWLEDGMENTS

B.G. acknowledges Dr. D.S. Kothari postdoctoral fellowship for financial support. We acknowledge Sahasrat, SERC, and TUE-CMS, SSCU at IISc, Bangalore, India for the computational facilities.

## MATERIALS AND METHODS

### Model Building

#### gp41 trimer

The modeled structure of ecto-transmembrane domain of HIV-1 gp41 post-fusion (PoF) six-helix bundled (6HB) conformer was borrowed from our previous work (33). In order to fit the gp41 trimer in the current work, we splayed the N-terminal FP and TMD of PoF structure in the opposite direction (Fig. S1B). The Martini coarse-grained (CG) model of the customized atomistic model of the gp41 trimer was obtained from CHARMM-GUI web portal (34, 35).

#### Human and HIV-1 cell membranes

A bilayer (dimension 40 × 40 nm^2^) and a vesicle (diameter ~20 nm) composed of mixed CG lipid types mimicking near biological compositions of the human T-cell membrane lipidome (11, 36) were constructed using Martini builder at CHARMM-GUI web interface. Similarly, a vesicle (diameter ~20 nm) with near biological lipid composition of HIV-1 cell membrane (11, 36, 37) was generated using Martini builder at CHARMM-GUI web interface. The inner and outer layers of cell membrane models were asymmetrically composed of eight different types of lipids (Table S1 in SM). The HIV-1 vesicle generated by CHARMM-GUI server contains 6 holes to allow passage of water and flip-flop of lipids between leaflets. The holes were gradually closed during equilibration stages and further simulated for 5*μ*s. The stable HIV-1 vesicle after the production run, with limited water, was used for the fusion study in this study. For more details see S11.

#### Human—gp41—HIV-1 complex

To study the gp41 mediated fusion mechanism of HIV-1 with human cell, we bridged the HIV-1 and human cell membrane CG models by gp41 trimers. The minimal distance between the HIV-1 and the human cell membranes were maintained ~8 Å initially. The TMD and FP domains of gp41 trimer were inserted in the HIV-1 and the human membrane models, respectively. The lipids of the membranes within 4 Å of the FP and TMD of the gp41 were deleted to remove any bad contacts. We build different human–gp41–HIV-1 complexes, where the count of gp41 trimers varies from zero to four. The ternary complex of the human membrane, gp41 trimer, and HIV-1 vesicle were placed in a box with a dimension of 40 × 40 × 36 nm3. The box with the complex was solvated by standard Martini water beads and 0.14 M NaCl salt.

### Simulation Details

Simulations were performed using Gromacs version 5.1.4 (38) with standard Martini v2.2 force field for protein, lipids, water, and ions (39). The solvated and neutralized systems were initially minimized by steepest descent algorithm with a force tolerance of 100 kJ/mol/nm. Equilibration of the systems were achieved by multiple stages, during which restraints applied on the lipids were gradually released and integration time steps were steadily increased from 10 fs to 30 fs. All the simulations were performed with periodic boundary conditions (PBC) to simulate bulk behavior. The dielectric constant was set to 15. The long range electrostatic Coulumbic interactions were truncated with a shift cutoff of 1.2 nm and the Lennard-Jones (LJ) potential was smoothly shifted to zero within 0.9 to 1.2 nm during CGMDS in this study. The pressure was maintained at 1 bar using Berendsen barostat during equilibration and production simulations. Semi-isotropic pressure coupling was adapted for all the systems, except for vesicle systems we applied isotropic coupling. The temperature was adjusted using Berendsen and V-rescale thermostats (40) during equilibration and production simulations, respectively. The gp41, HIV-1 vesicle, human bilayer, and solvent were coupled separately with a time constant of 1 ps. The neighbor list was updated at every 5 steps during the CGMDS with 30 fs time step.

#### CGMDS with varied number of gp41 trimers anchored

The simulation of solvated systems containing only the HIV-1 vesicle and human membrane models (vesicle and bilayer), without gp41 trimers, were simulated for 5 *μ*s each at 300 K. Further, the systems containing HIV-1 vesicle, human bilayer and bridged HIV-1 gp41 trimers (count varies from one to four) were simulated for 10 *μ*s each at 300 K.

#### CGMDS at varying temperatures

Simulation of systems containing vesicle, bilayer, and four HIV-1 gp41 trimers were further simulated at five different temperatures, viz. 300 K, 320 K, 340 K, 360 K, and 400 K. Each simulation was performed for 2 *μ*s and repeated twice with different starting velocity.

#### CGMDS of POPC/POPS bilayer-vesicle with four gp41 trimers at varying temperatures

Simulation of the bilayer and a vesicle composed of POPC (80%) and POPS (20%) lipids with four gp41 trimers were performed at 300 and 400 K for 10 *μ*s each. All simulations in the current study are bereft of any external force, electric field, membrane tension, and biased lipid compositions.

### Data Analysis

All the analyses were carried out by using home-written codes and/or the tools available in GROMACS package. Movies and snapshots were made using Visual Molecular Dynamics software (41). Any lipid from human bilayer moving more than 15 Å above from the geometrical center of FP and TMD domain of 4 gp41 trimers are considered to be transferred from human cell membrane to HIV-1 vesicle (C→H). Similarly, any lipid from the vesicle moving more than 15 Å below from the geometrical center of FP and TMD domain of 4 gp41 trimers are considered to be transferred from HIV-1 vesicle to human cell membrane (H→C).

