## Supplementary files for "Concerted interactions between multiple gp41 trimers and the target cell lipidome may be required for HIV-1 entry"

**This PDF file includes:**
Figs. S1 to S11
Table S1

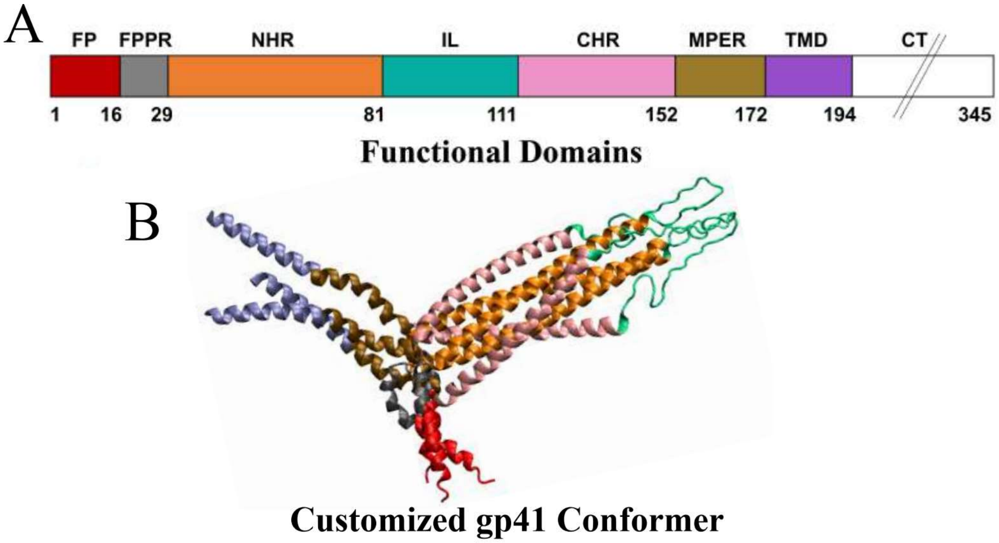


**Fig. S1. The functional domains of HIV-1 gp41.** *(A)* Different colored bars are used to represent fusion peptide (FP), fusion peptide proximal region (FPPR), N-terminal heptad repeat (NHR), immunodominant loop (IL), C-terminal heptad repeat (CHR), membrane-proximal external region (MPER), transmembrane domain (TMD), and cytoplasmic domain (CD). The cartoon representaion of trimer of the HIV-1 gp41 TMD-ectodomain region (residues 1–194), used in the current study, is shown. The conformer is generated after splaying the TMD and FP of modeled post fusion conformer of gp41 trimer, taken from our previous study, in opposite direction.

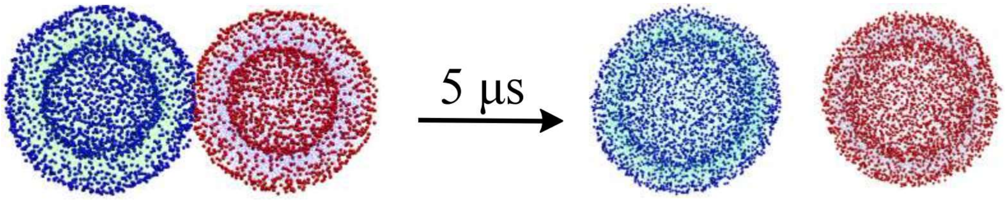


**Fig. S2. Simulation of the human and HIV-1 vesicle membrane models, with near biological compositions, in the absence of gp41.** The initial (left) and final (right) configurations of the simulated system at 300 K containing the human and HIV-1 vesicles without any gp41 trimer are shown. The lipid head groups and tails of the human vesicle are shown in vdW (blue) and line (green) representations, respectively. The lipid head groups and tails of the HIV-1 vesicle are shown in vdW (red) and line (light blue) representations, respectively. Water and ions are not shown here for clarity.

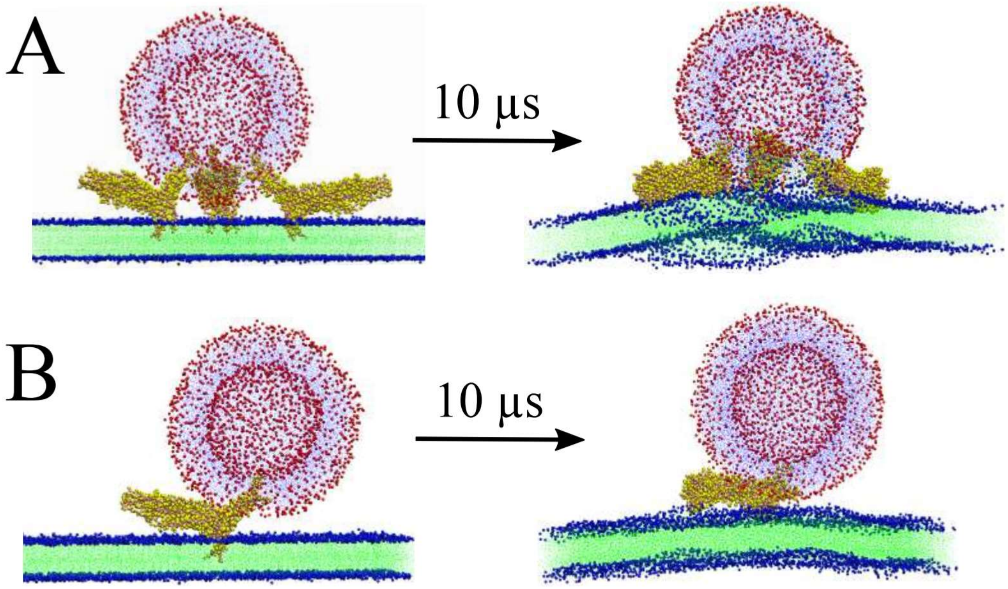


**Fig. S3. Simulation of the human and HIV-1 membrane models, with near biological compositions, in the presence of gp41.** The initial (left) and final (right) configurations of the simulated system at 300 K consisting of the human bilayer and HIV-1 vesicle in the presence of (*A*) triple and (*B*) single gp41 trimeric units are depicted. The lipid head groups and tails of the human bilayer are shown in vdW (blue) and line (green) representations, respectively. The lipid head groups and tails of the HIV-1 vesicle are shown in vdW (red) and line (light blue) representations, respectively. The gp41 trimeric unit is shown in vdW representation with yellow color. Water and ions are not shown here for clarity.

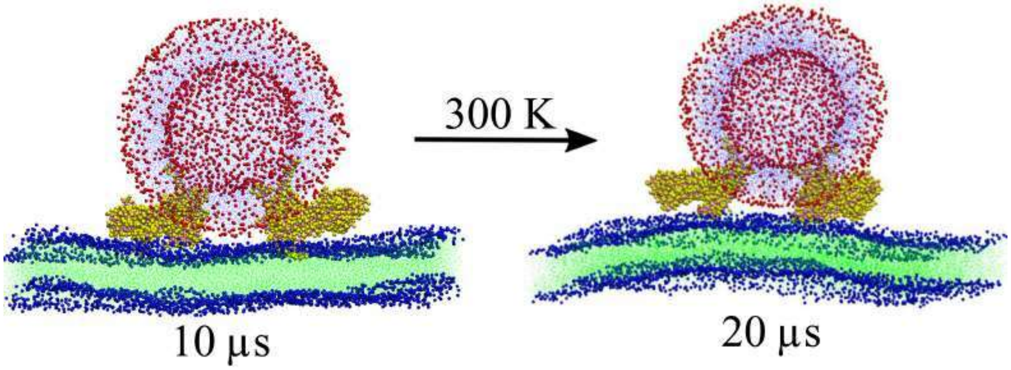


**Fig. S4. Extension of simulation of the human and HIV-1 membrane models in the presence of 2 gp41 trimers.** The configurations of the simulated system at 300 K at 10 $\mu$s (left) and at 20 $\mu$s (right) consisting of the human bilayer and HIV-1 vesicle in the presence of double gp41 trimeric units are depicted. The lipid head groups and tails of the human bilayer are shown in vdW (blue) and line (green) representations, respectively. The lipid head groups and tails of the HIV-1 vesicle are shown in vdW (red) and line (light blue) representations, respectively. The gp41 trimeric unit is shown in vdW representation with yellow color. Water and ions are not shown here for clarity.

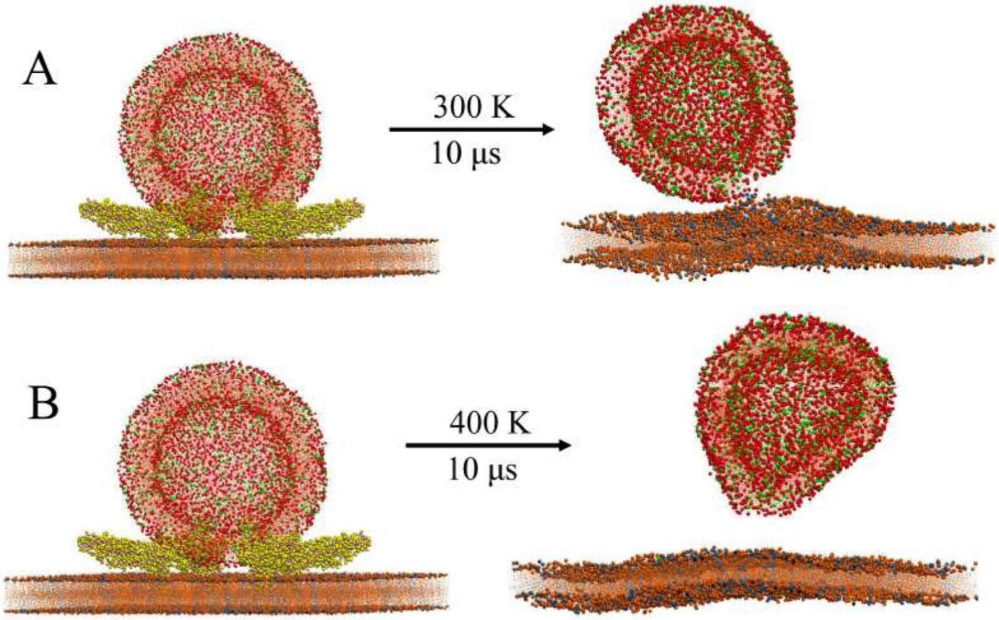


**Fig. S5. Simulation of the human and HIV-1 membrane models, with near arbitrary lipid compositions, in the presence of gp41.** The snapshots of the initial (left) and final (right) configurations of the human bilayer and HIV-1 vesicle composed of POPC and POPS in 80:20 proportion at (*A*) 300 K, and (*B*) 400 K for 10 $\mu$s are shown. The POPC and POPS lipids of the human bilayer are shown in brown and blue colors, respectively. The POPC and POPS lipids of the HIV-1 vesicle are shown in red and green colors, respectively. The gp41 trimer is depicted in yellow vdW representations. The exchange of lipids between human and HIV-1 cell membranes with non-biological compositions at room temperature, as well as elevated temperatures are not observed. This observation justifies the complex compostion adapted by the cell membranes to accomplish the critical biological phenomenons. The gp41 in the final configuration, water and ions are not shown here for clarity.

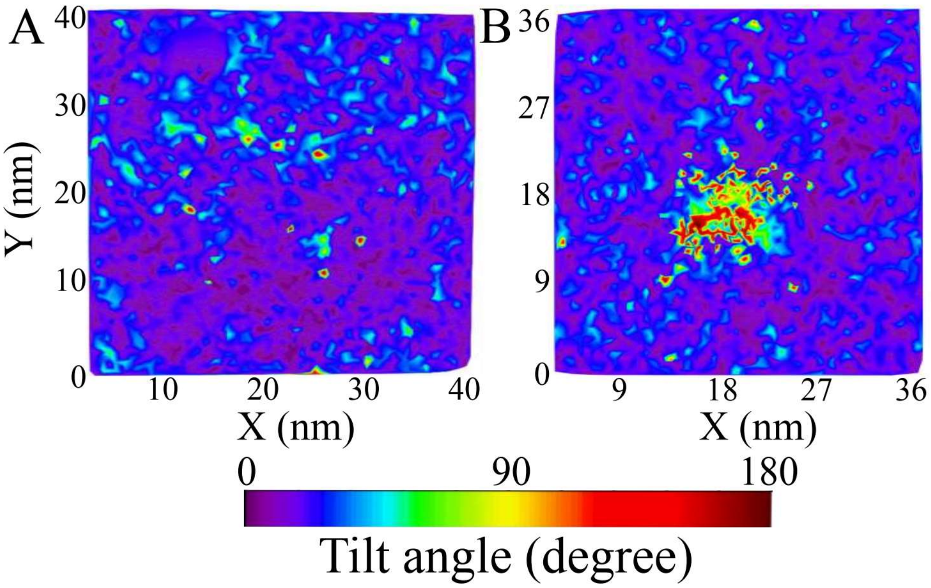


**Fig. S6. Lipid tilt distribution map.** The surface distribution of tilt angle of lipids of the human bilayer at (*A*) the beginning of simulation, and (*B*) at beginning of stalk formation is shown.

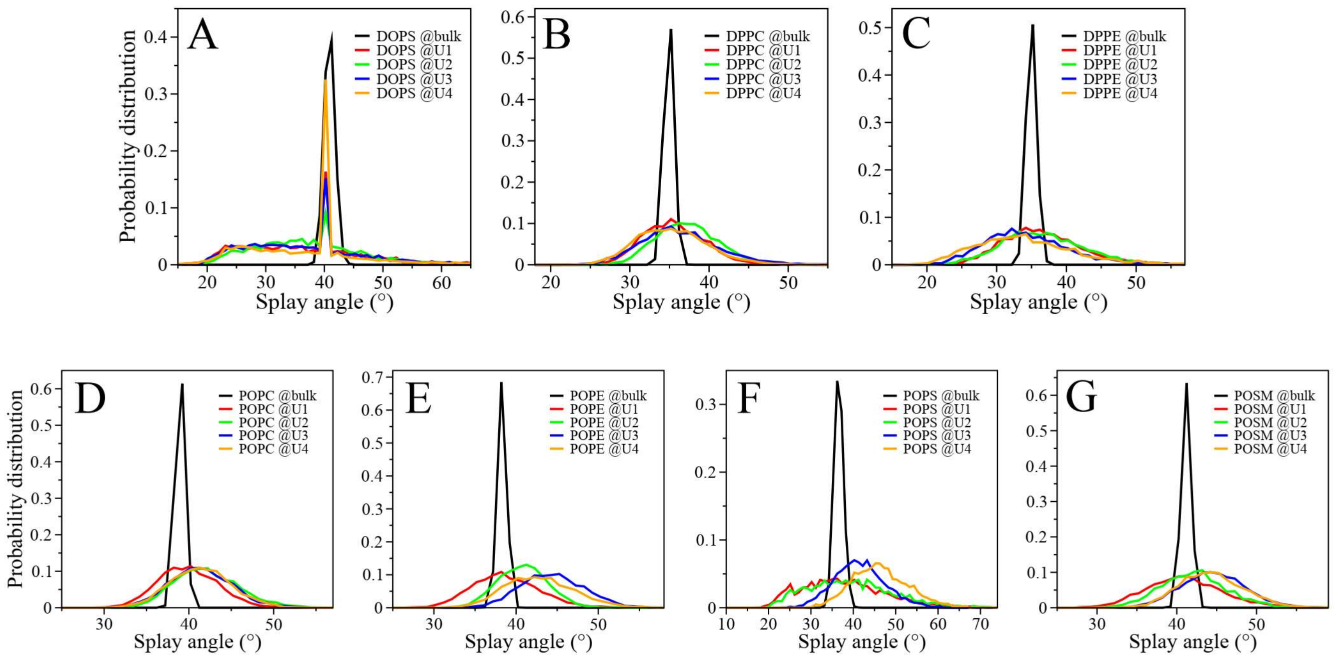


**Fig. S7. Splay angle probability distribution.** The normalized probability distribution of splay angle between the two tails of (*A*) DOPS, (*B*) DPPC, (*C*) DPPE, (*D*) POPC, (*E*) POPE, (*F*) POPS, and (*G*) POSM lipids at bulk (black line), and lipids around gp41 (20 Å of center of mass of FP+TMD domain) of each trimeric units (U1-red line, U2-green line, U3-blue line, and U4-orange line) is shown.

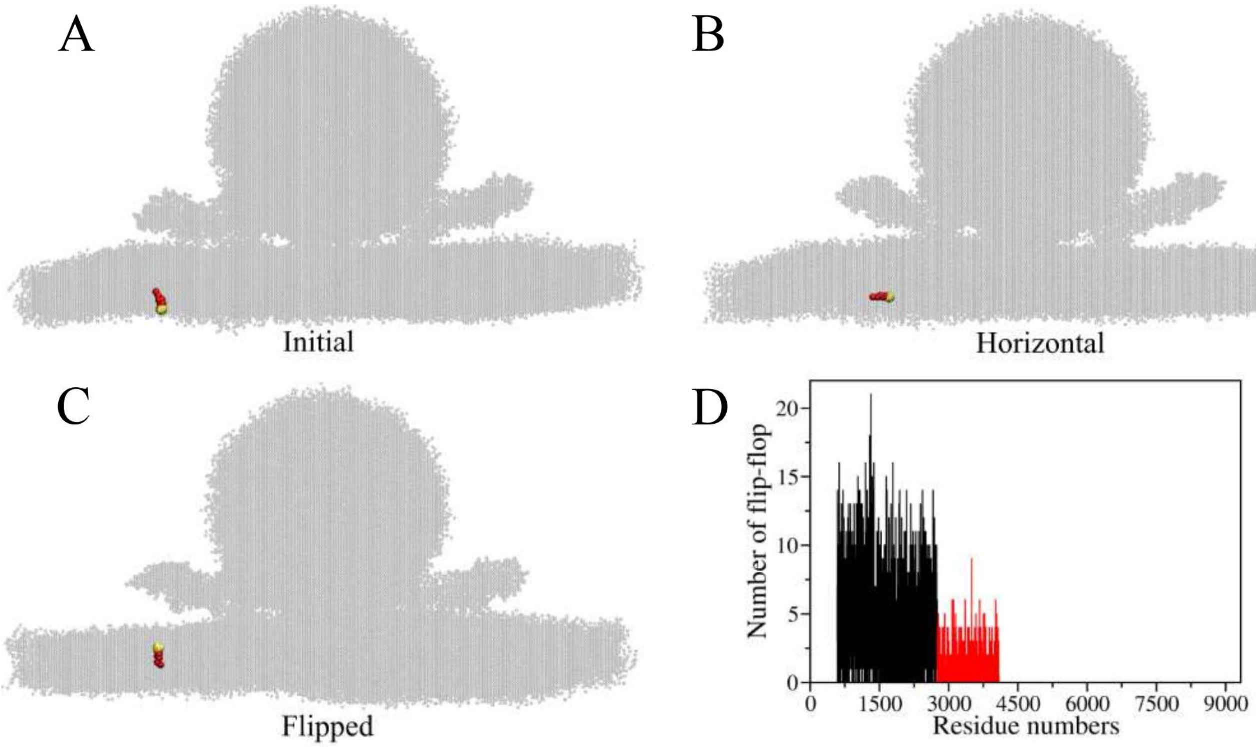


**Fig. S8. Flip-flop of lipids.** Snapshots displaying the major events during the flip-flop pathway of cholesterol: (*A*) initial stage: cholesterol at the lower leaflet, (*B*) intermediate stage: horizontal alignmnet of cholesterol, and (*C*) final stage: cholesterol move to the upper leaflet. Hydroxyl head group and non-polar fatty acid tail of cholesterol is represented by yellow and red vdW spheres, respectively. (*D*) The number of times each lipid flipped between the leaflets, before stalk formation, are shown in the plot. We noticed that only cholesterols (residue number 583–4084) underwent translocation within the leaflets and the rate of translocation between leaflets is higher for cholesterols of human bilayer (black: residue number 583–2749) than cholesterols of HIV-1 vesicle (red: residue number 2750–4084).

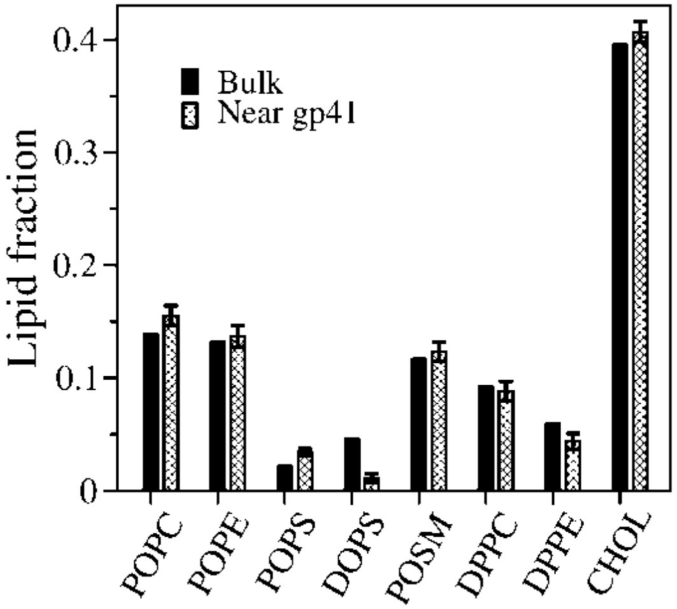


**Fig. S9. Histogram plot of lipid fractions.** Fraction of each lipids at the bulk (not within 20 Å of center of mass of gp41 FP+TMD domain) and near gp41 trimeric units (within 20 Å of center of mass of gp41 FP+TMD domain) calculated during the simulation time is shown.

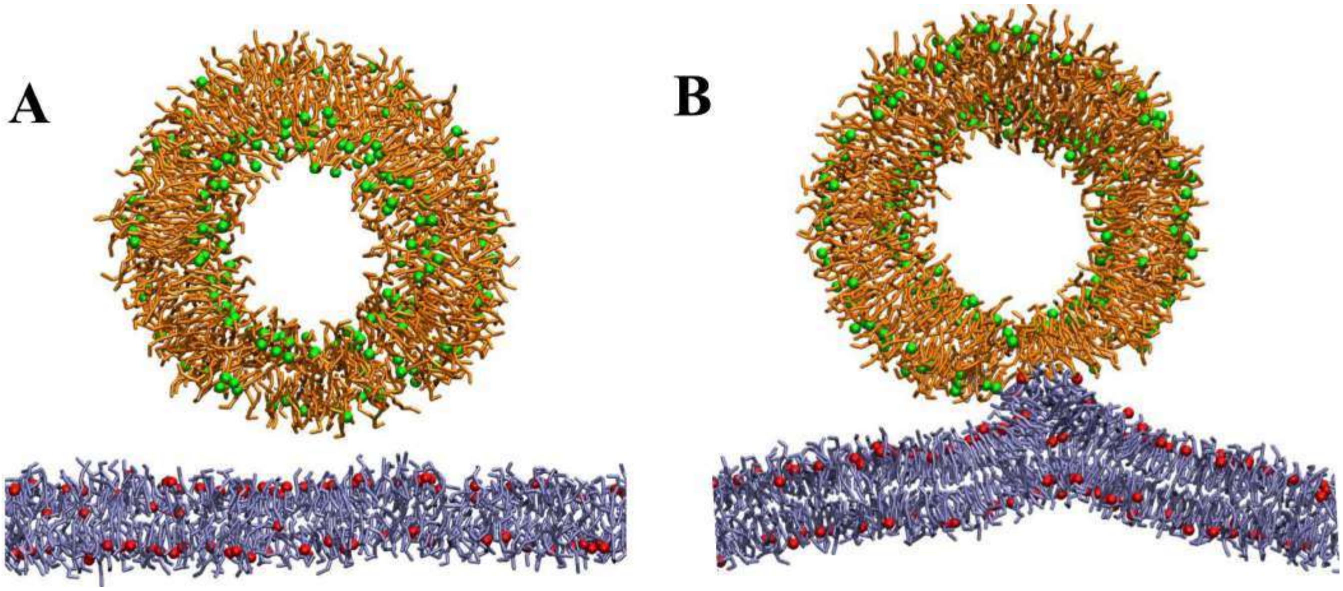


**Fig. S10. Stalk initiation snapshot.** The clipped image of the (*A*) initial and (*B*) stalk initiation stage during simulation of HIV-1 and human bilayer using 4 gp41 trimers at 300 K is depicted. The HIV-1 and human membrane models are represented by vesicle and bilayer, respectively. Water, ions and gp41 units are not shown here for the clarity.

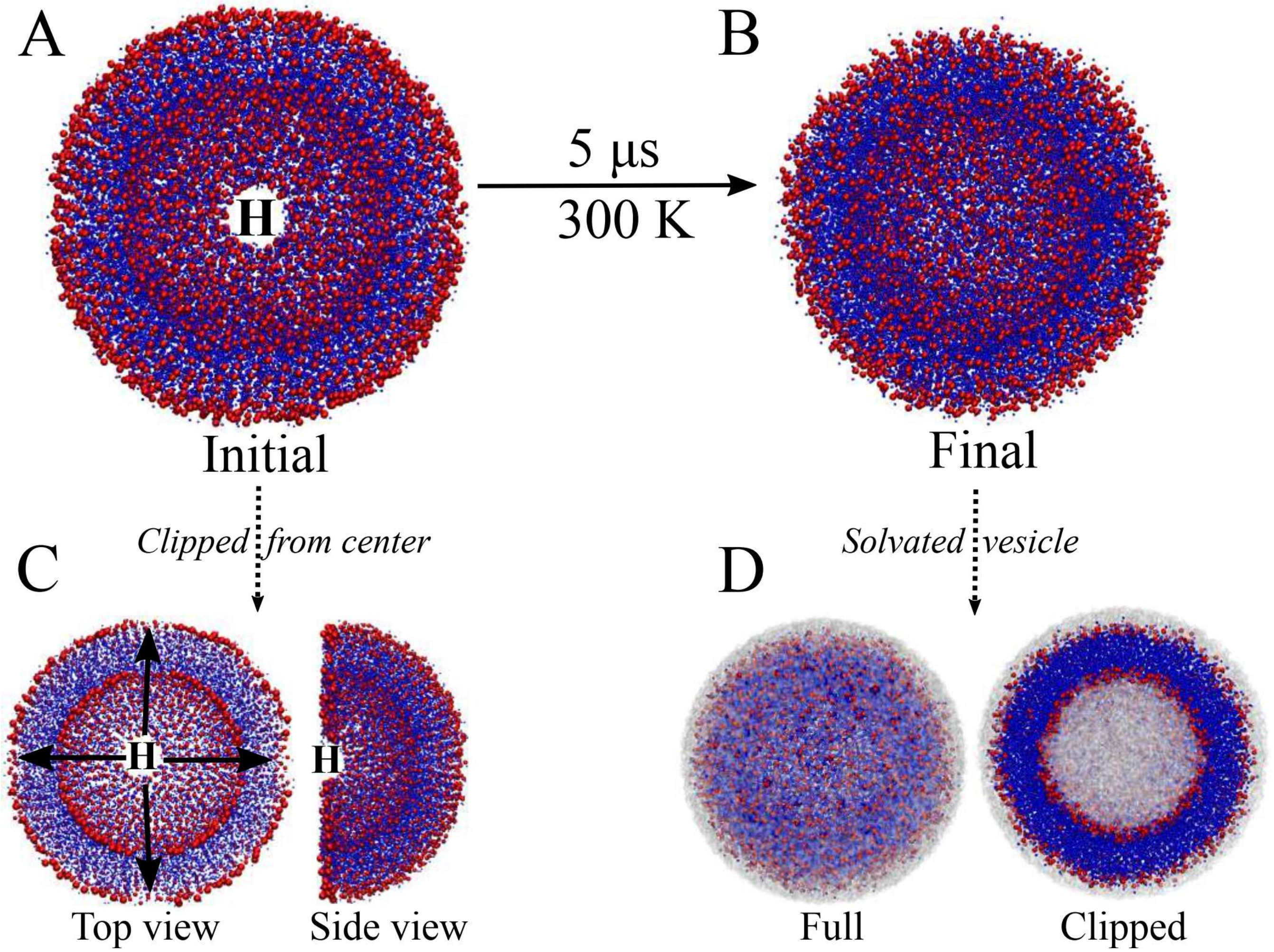


**Fig. S11. Equilibration of HIV-1 vesicle.** Initially, the HIV-1 vesicle obtained from CHARMM-GUI web server contains (*A*) 6 holes (H). (*C*) The location of holes are shown in the top and side view of HIV-1 vesicle, clipped from the center. The initial structure was subjected to CGMD equilibration and production run to obtain a (*B*) stable vesicle without holes. The stable HIV-1 vesicle, with (*D*) a water layer (surface representation) of around 11 nm radius from the center of the vesicle is used for further fusion study. The lipid head groups and tails of the HIV-1 vesicle are shown in vdW (red) and dots ( blue) representations, respectively. The simulation was performed in the presence of ions and explicit coarse-grained water model, but not shown in (*A*) and (*B*) for clarity.

**Table S1. The lipid composition of HIV-1 cell membrane and human T-cell membrane.**

| Lipid | HIV-1 | | Human | |
|---|---|---|---|---|
| | IL | OL | IL | OL |
| 1. DPPC | 0 | 0 | 208 | 591 |
| 2. DPPE | 0 | 0 | 176 | 335 |
| 3. DOPS | 0 | 0 | 389 | 0 |
| 4. POPC | 64 | 223 | 643 | 279 |
| 5. POPE | 313 | 211 | 155 | 460 |
| 6. POPS | 153 | 34 | 0 | 0 |
| 7. POSM | 66 | 524 | 306 | 117 |
| 8. CHOL | 609 | 726 | 1079 | 1088 |

The membranes are composed by dipalmitoyl phosphatidylcholine (DPPC), dipalmitoyl phosphatidylethanolamine (DPPE), dioleoyl phosphoserine (DOPS), palmitoyl oleoyl phosphocholine (POPC), palmitoyl oleoyl phosphatidylethanolamine (POPE), palmitoyl oleoyl phosphoserine (POPS) palmitoyl oleoyl sphingomyelin (POSM), and cholesterol (CHOL). The lipid composition of HIV-1 and human T-cell membranes are highly inhomogeneous, with asymmetric inner leaflet (IL) and outer leaflet (OL) lipid content.
